# When Mechanical Stress Matters: Generation of Polyploid Giant Cancer Cells in Tumor-like Microcapsules

**DOI:** 10.1101/2022.09.22.508846

**Authors:** Adrian Bühler, René Krüger, Mahshid Monavari, Miguel Fuentes-Chandía, Ralf Palmisano, Johannes Schödel, Aldo R. Boccaccini, Anja K. Boßerhoff, Melanie Kappelmann-Fenzl, Gaelle Letort, Aldo Leal-Egaña

**Author notes:** Corresponding author. Aldo Leal-Egaña. These authors contributed equally to this work (Adrian Bühler, René Krüger, Mahshid Monavari).

## Abstract

In this work, we studied the generation and rising of polyploid cancer cells as a product of mechanical stress. To this purpose, MCF7 breast cancer cells were cultured on 2D (*i.e*. flasks, or flat hydrogels), and in 3D milieus (*i.e*. Spheroids, or immobilized within alginate-gelatin microbeads, named in this work as tumor-like microcapsules), and further analyzed by biophysical and genetic methods (i.e. single-cell Traction Force Microscopy and RNA-seq respectively).

Our results show that MCF7 cells preconditioned onto 2D surfaces exhibit a low number of polynucleated cells, while their culture in 3D environments triggered their progressive generation with time. Genetic studies enabled us to determine that polyploid cells found in tumor-like microcapsules are likely originated by cell-cell fusion and disrupted cytokinesis, showing most of the genetic markers for Polyploid Giant Cancer Cell, while cells cultured as spheroids seem to be likely generated by other mechanisms, such as cell cannibalisms, entosis, or emperipolesis.

Our outcomes strongly suggest that both mechanical stress and confinement are required to stimulate cell polyploidy, which can be easily addressed by the immobilization of breast cancer cells in tumor-like microcapsules.

## 1. Introduction

Intratumor heterogeneity describes the existence of neoplastic tissues constituted by a wide variety of malignant cells, which are originated from one single malignant clone. Heterogeneous populations are traditionally co-localized in the same milieu, exhibiting distinct biophysical, physiological or genetic features, varying from cell-to-cell.^[1–4]^

Within heterogeneous cell types already described in breast cancer pathologies, polynucleated cells have attracted the attention of researchers working in this field.^[5–10]^ These are characterized by their larger spreading area and higher number of nuclei with respect to healthy cells, or even to other cancer cells found in the same milieu.^[5–17]^ As already reported *in vivo,* polynucleated cells can spontaneously develop inside tumors by different mechanisms, including cell-in-cell growth or cannibalism (also known as cell-eat-cell),^[11–17]^ as well as by cell-cell fusion or disrupted cytokinesis.^[12,3,5–17]^ These mechanisms mostly differ in the way how polynucleated cells are generated: while cell-in-cell or cell-eat-cell are characterized by the existence of a cell endocyted by another one (enabling their growth or even proliferation inside the host cell), the processes of cell-cell fusion or incomplete cytokinesis are mostly mediated by membrane interactions and cytoskeleton failures, merging two independent and/or colocalized cells, existing no host.^[11,17]^

Clinically speaking, polyploid cells seem to be involved in stemness, development of resistance to chemotherapeutic treatments, metastasis to bone, as well as tumor relapse.^[2–19]^ Hence, efforts have been made on searching for alternatives to generate polyploid cancer cells at laboratory scale.^[14–17]^ In particular, the type known as Polyploid

Giant Cancer Cells (PGCC) has been widely studied, and protocols to produce them *in vitro* have been established. For example, based on their capability to disrupt the cell cycle, the use of several drugs -such as Paclitaxel, Doxorubicin or Docetaxel-has been widely employed. Additionally, CoCl_2_, radiation (*i.e*. from 6 to 14 Gy), or even hyperthermia (*i.e*. 42°C) have been also reported as traditional strategy to stimulate their rising.^[14–17]^ However, it is important to note that none of these stimuli properly resemble the conditions sensed by cancer cells located inside primary tumors.

Thus, with the purpose to stimulate the formation of polynucleated cells in milieus resembling the biophysical properties already described for malignant tissues, other strategies have been tested. For example, new studies have shown that cell entosis can be triggered by anchorage independence and glucose starvation.^[20]^ Additionally, the generation of polynucleated cells have been also observed when cancer cells are cultured in presence of nanoparticles, or stimulated by the use of new anticancer molecules.^[21,22]^ However, since these methods are mostly focused on describing the existence and morphological characteristics of multinucleated cells, there is a lack of information regarding their generation and rising in time.

To this purpose, the use of 3D polymer-based matrices, mimicking the most relevant properties of neoplastic niches, has emerged as a good alternative to fulfil this gap. In this regard, we recently published that the increased resistance of encapsulated MCF7 cancer cells to cisplatin,^[28,29]^ seems to be due to the enhanced presence of polynucleated cells found in alginate-gelatin microcapsules (also known as tumor-like microcapsules in this manuscript).^[28,29]^ However, the way how these polynucleated cells are increasing in number in absence of any external stimuli, and which 3D experimental system may exhibit advantages to study polyploidy *in vitro,* are still unsolved questions.

Thus, in this work we report the results obtained after culturing the breast cancer cell line MCF7 in different two-dimensional (2D) and three-dimensional (3D) milieus. These platforms, in combination to time-lapse assays, would offer the possibility to analyze the generation of polyploid cells *in vitro,* under experimental conditions similar to those found in the neoplastic milieu.

## 2. Materials and Methods

### 2.1 Cell culture

The breast cancer cell line MCF7 (DSMZ Cat. N° ACC 115) was sourced from our departmental cell bank, and were cultured in DMEM cell culture media (High glucose, Gibco, USA), supplemented with 10% v/v of FBS (Fetal Bovine Serum; Corning, USA) and 1.0% v/v penicillin-streptomycin (Gibco, USA).

### 2.2 Cell culture on 2D: flat surfaces and flat hydrogels

MCF7 cells were cultured on cell culture flasks (175 cm^2^, Corning), keeping their confluency between 30% and 80%. Flat hydrogels were made of polyacrylamide functionalized with collagen type I (Sigma-Aldrich) and bound to (bound to 22 × 22 mm coverslips). These gels were made following protocols previously Tse et al.^[30]^

Flat surfaces (either hard or soft) were used to culture cancer cells for 5 days prior to analysis. When characterizing them by single-cell traction force microscopy, a number of 10·10^3^ cells/cm^2^ were seeded.

### 2.3 Preparation of 3D suspended spheroids and tumor-like microcapsules

Suspended spheroids were obtained by seeding 20·10^3^ cells in 24 well plates coated with 250 μl of agarose (1.0% w/v dissolved in sterile PBS). This assay was carried out according to previously published methods.^[20,21]^

Tumor-like microcapsules were made using a using a micro-encapsulator device Büchi-390, according to methods previously published.^[28,29]^ Microcapsules were constituted of 1.0% w/v alginic acid (PH163S2, JRS Pharma GmbH, Germany) and 1.0% w/v gelatin type A (Sigma-Aldrich, Germany). These polymers were dissolved in a solution containing 0.1M HEPES (pH 7.4, Sigma-Aldrich, Germany), 1.0 % w/v NaCl and 1.0 % v/v penicillinstreptomycin (Sigma-Aldrich, Germany). This blend was then filtered by using a 0.45 μm pore diameter filter (Merck Millipore, Germany), and loaded with MCF7 cells with a density of 5·10^5^ cells/mL.

To crosslink the alginate-gelatin microcapsules, a solution of 0.6 M of CaCl_2_ (dissolved in 1.0% w/v HEPES and containing 1.0% w/v NaCl; pH 7.4) was used.

### 2.4 Determination of the elasticity of alginate-gelatin microcapsules

The stiffness of tumor-like microcapsules was determined by nanoindentation. To this purpose, a nano-indenter PIUMA (Optics 11, Netherlands) was used. To this purpose, alginate-gelatin microbeads of 1.5 mm of diameter were made and placed in an 3D printed reservoir (filled with cell culture media), to avoid their rolling or displacement. The elasticity of the microcapsules was determined immediately after encapsulation, as well as after 3, 6 and 10 days post-encapsulation.

### 2.5 Single cell traction force microscopy assays

Single Cell Traction Force Microscopy (TFM) assays were performed according to published methods.^[31,32]^ Briefly, cells were seeded on flat polyacrylamide gels coated with collagen type-I. These hydrogels had an elasticity of 8.5 ± 2.8 kPa. Forces exerted by single cells were determined by computational analysis using a software gently sourced by Dr. Martial Balland, Laboratoire Interdisciplinaire de Physique, Grenoble, France.

### 2.6 Cell imaging

Cells were imaged using a Spinning Disc Axio Observer Z1 from Zeiss, placed at the Optical Imaging Centre Erlangen (OICE). For determining the existence of polynucleated cells, MCF7 were anchored to glass coverslips (22 × 22 mm), and fixed using a 4.0% w/v paraformaldehyde solution (Sigma Aldrich, USA). This step was followed by their staining with Phalloidin-Rhodamine (Invitrogen, Germany), according to the provider’s instructions. The spreading area of anchored cells was measured and analyzed with the open-source Fiji software. Statistical analyses were performed with the open-source software R.

### 2.7 Cell proliferation assays

Studies of cell duplication were carried out with cells cultured in 3D using published methods.^[33]^ Briefly, around microcapsules and spheroids were placed in a 48 well plate. These were cultured in presence of WST-8 (10% v/v, Thermofisher, Germany), for three hours, before we carried out the measurements, in accordance with the provider instructions. After measuring the metabolization of WST-8, capsules and spheroids were washed with a 1.0% w/v HEPES solution (containing 1.0% w/v NaCl; pH 7.4), and kept in culture with cell culture media. This strategy was repeated during 10 days with the same samples. Proliferation assays were carried out in 5 replicas.

### 2.8 Isolation of cells from alginate-gelatin microcapsules

Cells were isolated from the microcapsules by immersing them in a solution containing of 22.5 mM of sodium citrate dihydrate, 60 mM EDTA and 150 mM of sodium chloride (Sigma-Aldrich; pH 7.4) for 1 minute, following protocols previously published.^[28,29]^ After that, cells were isolated by centrifugation (1300 rpm, 4 min).

### 2.9 RNA-Seq assays and transcriptome analysis

RNA extraction was carried out according to protocols already published.^[28,29]^ All experiments were carried out in triplicate. 75 bps paired-end sequencing was performed on a HighSeq 4000 Sequencer (Illumina) with an average number of 20 million reads per sample. After a quality check using FastQC v0.11.9,^[34]^ reads were aligned to the human reference genome (hg38) using the STAR alignment software (v 2.5.2a).^[35]^ After mapping, only reads that mapped to a single unique location were considered for further analysis. The mapped reads were then used to generate a count table using the feature-counts software (v 1.4.6-p5).^[36]^ The raw reads were filtered, normalized, and visualized by using R version 4.0.1. The DESeq2 package, version 1.26.0, was used for logarithmic transformation of the data and for data exploration. The differential expression analysis was done using the DESeq2 standard approach. Adjusted p-values are calculated using the Benjamini–Hochberg method within DESeq2. Differentially expressed genes with a log2FC+/-1.5 and p-adjusted <0.01 were regarded as statistically significant. Gene annotations were using biomaRt version 2.50.3.^[37]^

The different expression of the immobilized cells v/s suspended spheroids was performed and the Reads Per Kilo-base of transcript per Million mapped reads (RPKM) was calculated. RPKM was used to normalize the raw reads. Gene annotations were added to the RPKM using biomaRt version 2.50.3.^[38]^. Subsequently, we focused on comparing genes traditionally described for characterizing the presence of PGCC and cell-eat-cell. These genes are listed in the Supplementary Table S2. These genes were then clustered with the complete linkage method, and plotted using the pheatmap package version 1.0.12.^[39]^

#### 2.9.1 Cluster profile assays

Functional enrichment analysis of differential expressed genes, of the immobilized cells versus suspended spheroids, was performed.^[40]^ Functional data analysis was performed using the Gene Set Enrichment Analysis (GSEA) GSEAPreranked tool (GSEA; v4.1.0). GSEA analysis was performed using preranked log2FC with an adjusted p-value < 0.01 from DEG analysis. The analysis using C5 GO biological processes gene sets from MsigDB (v7.5.1) was performed with 1000 permutations of gene sets.^[41]^ Enrichment results were considered significant with FDR < 0.25 and p-value <0.05. Corresponding Enrichment Maps were created by the Cytoscape Software Tool (v3.9.1). AutoAnnotate with the Markov CLustering Algorithm (MCL) was used to identify clusters and visually annotating them (v1.3.5).^[42]^

### 2.10 Statistical analysis

To determine the reference values of Total Forces v/s Area (square), we took 0.05 and 0.95 quantiles of the values exhibited by the whole analyzed population of cancer cells, in terms of tractions forces or cell area. This obtained rectangle has dimensions of 21.6 × 87.2 nN in forces (i.e. 90% of control cells exhibit their traction forces within this range) and 299.2 × 627.2 μm^2^ (i.e. 90% of control cells display their spread area within these values). We used this rectangle as a way to determine the morpho-mechanical differences between cells cultured on flat surfaces, and cells cultured on other different 2D or 3D culture conditions. We compared the distribution of the Force/Area values after 1, 2, 4, 7 and 10 days for cells cultured in ether microcapsules or spheroids to the Force/Area values for cells cultured on cell culture flasks (2D control) with the Kolmogorov-Smirnov test. All statistical analysis were performed within R. In each case, the minimum number of analyzed cells was 65.

## 3. Results and Discussions

In breast pathologies, tumors are characterized by their capability to entrap malignant cells, isolating them from healthy tissues and organs.^[23,42]^ Due to the increased presence of catalytic enzymes crosslinking collagen fibers (i.e. Lysyl Oxidase), as well as the secretion of Tissue Inhibitors of Metalloproteinases by Cancer Associated Fibroblasts, during the progression of this pathology tumors enhance their elasticity from ~4kPa (at initial stages of this pathology), up to ~80 kPa (in advanced phases of this disease).^[43]^ The effect of the cell confinement in stiffer 3D environments triggers the expression of biophysical and genetic hallmarks, which varies from cell-to-cell, leading the generation of heterogeneous populations,^[1,3,17,23,44]^ Therefore, in this work we hypothesize that confinement and mechanical stress may trigger the generation of polynucleated cells *in vitro*.

To test this, cells were pre-conditioned to different 2D and 3D milieus (Figure 1A), before being analyzed by biophysical and genetic methods. 2D surfaces represent the most traditional platforms employed to culture cancer cells, while 3D milieus exemplify the new technical alternatives used to study the behavior of malignant cells *in vitro*.^[23,45–47]^ Concerning the use of soft 2D hydrogels to pre-condition cancer cells, since these have a similar elasticity to the tumor-like microcapsules (23 ±3.0 kPa or 21 ±2.0 kPa respectively; Figures 1B and 1C), this strategy was utilized to discern the respective impact of the matrix stiffness upon polyploidy (i.e. 2D v/s 3D). It is also important to mention that the tumor-like microcapsules share the mechanical properties to early primary breast tumors,^[15,25]^ representing an accurate model to study the influence of the elasticity and/or the confinement in cell polyploidization.

**Figure 1:**
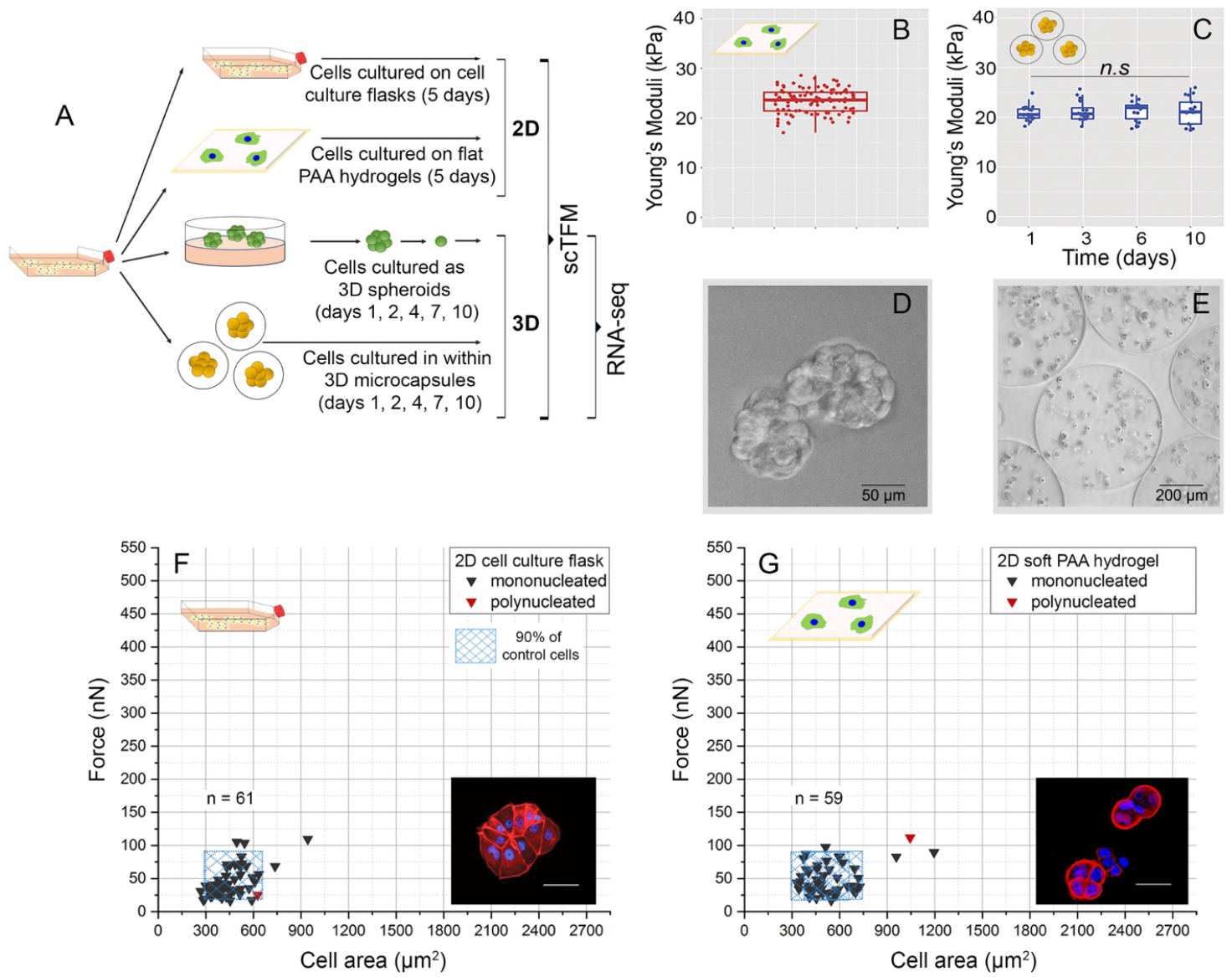
(A) Experimental design carried out in this work. MCF7 breast cancer cells were pre-conditioned onto 2D surfaces or in 3D milieus for 5 days. The elastic properties of 2D flat hydrogels (B) and the bulk of the alginate-gelatin microcapsules (C) show values of 23 ±3.0 kPa or 21 ±2.0 kPa respectively, measured as Young’s Moduli. Images D and E show the experimental systems to cultivate cells in 3D. Picture D shows cells forming spheroids, while image E corresponds to immobilized MCF7 in alginate-gelatin microcapsules. Plots F and G depict the morpho-mechanical characterization of MCF7 cells cultured on 2D surfaces, analyzed by single-cell Traction Force Microscopy. In these assays, cells were cultured on flasks (F) or on soft-hydrogels (G), and “n”represents the number of single cells analyzed. Mononucleated cells are represented as black triangles, and polynucleated as red ones. The blue rectangle encloses 90% of the measurements for each dimension of the cells cultured on flasks, which is considered the control experiment. The images located in (F) and (G) show the characteristic morphology of cells isolated from the cell culture flasks or cultured on soft hydrogels respectively. In these pictures, the red staining corresponds to the actin cytoskeleton, while the blue color depicts the presence of the nuclei. The scale bar corresponds to 25 μm.

Concerning the mechanical properties of spheroids, reported values for cell aggregates show that their elasticity resembles the stiffness of healthy tissues (approximately 6.0 kPa measured as Young’s Moduli).^[48–50]^ Images 1D and 1E show examples of the experimental systems used to culture MCF7 cells in 3D.

When using 2D surfaces, cells were cultured for 5 days before being isolated and analyzed. Initially, cells were characterized by single-cell Traction Force Microscopy (scTFM). This biophysical technique is aimed to determine the biomechanical activity of single cells, after being attached to substrates with controlled deformability.^[31,32]^ This information is relevant to determine the activity of cancer cells at different stages of the malignant progression (i.e. migration, invasion, metastasis), all processes during which neoplastic cells need to anchor, prior forming secondary tumors.^[32,51]^ Additionally, since this is an optical technique, this method enables comparing the anchorage surfaces of attached cells, as well as other features, such as the number of nuclei found in each cell.

As Figures 1F and 1G show, cells cultured on both 2D milieus exhibit a similar morpho-mechanical behavior, lacking of a relevant number of polyploid cells. To facilitate the comparison between all the experimental conditions, a square enclosing 90% of the measurements from the control experiment (*i.e*. cells cultured on flasks) was included in these plots. To determine the boundaries of this population, we took the 0.05 and 0.95 quantiles of cell areas and traction forces as limits, resulting in these dimensions: a range of 21.6 to 87.2 nN for force (i.e. 90% of control cells exhibit their traction forces within the defined range) and a range of 299.2 to 627.2 μm2 for spread area (i.e. 90% of control cells display their spread area within that range).

Concerning the results displayed in Figures 1F and 1G. it is important to mention that even though cells pre-conditioned onto culture plates are smaller (i.e. in terms of area), the exertion of forces is proportional to their spreading surface, keeping the Force/Area (F/A) ratio constant (Supplementary Information S1). These results demonstrate that flat surfaces (either hard or soft) are not playing a crucial role on polyploidization, probably having a stronger influence during cell migration (*i.e*. durotaxis).^[52]^

Besides studying the biophysical properties of cancer cells cultured on flat surfaces, MCF7 were cultured as spheroids or in tumor-like microcapsules, and further characterized by scTFM. In these experiments, samples were obtained after 1, 2, 4, 7 and 10 days post cell seeding/immobilization. As Figures 2 and 3 depict, cells isolated from both 3D systems enhanced their morpho-mechanical activity with respect to cells cultured on 2D, showing a stronger cell-matrix attachment and larger surfaces. Furthermore, and as shown in Figures 2 and 3, this behavior is dynamically enhanced with time. Additionally, in both 3D systems is possible to detect the apparition of polynucleated cells, a process which is more likely observed in the population extracted from the polymer-based microcapsules, instead of in cells isolated from spheroids.

**Figure 2:**
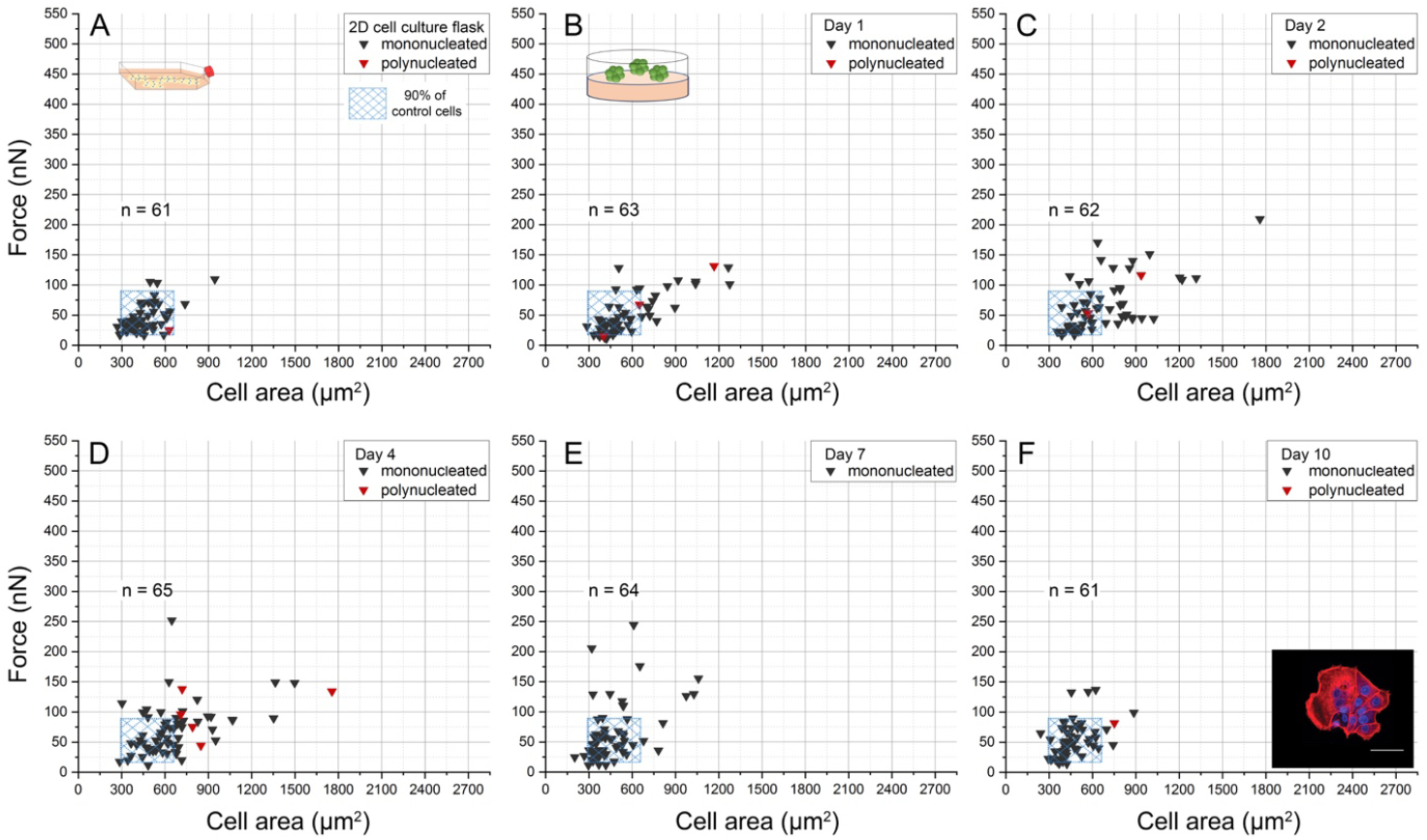
Morpho-mechanical characterization of MCF7 cells cultured on flasks for 5 days (A) or those isolated from suspended spheroids (B-F). In the last case, cells were isolated at days 1 (B), 2 (C), 4 (D), 7 (E) and 10 (F) post seeding. In all plots, the blue rectangle represents 90% of the measurements for each dimension of the cells cultured on flasks and is used as optic reference in each graphic. In all graphics, “n” represents the number of single cells analyzed. The image located in (F) shows the characteristic morphology of cells isolated from spheroids. In that picture, the red staining corresponds to the actin cytoskeleton, while the blue color depicts the presence of the nuclei. The scale bar corresponds to 25 μm.

**Figure 3:**
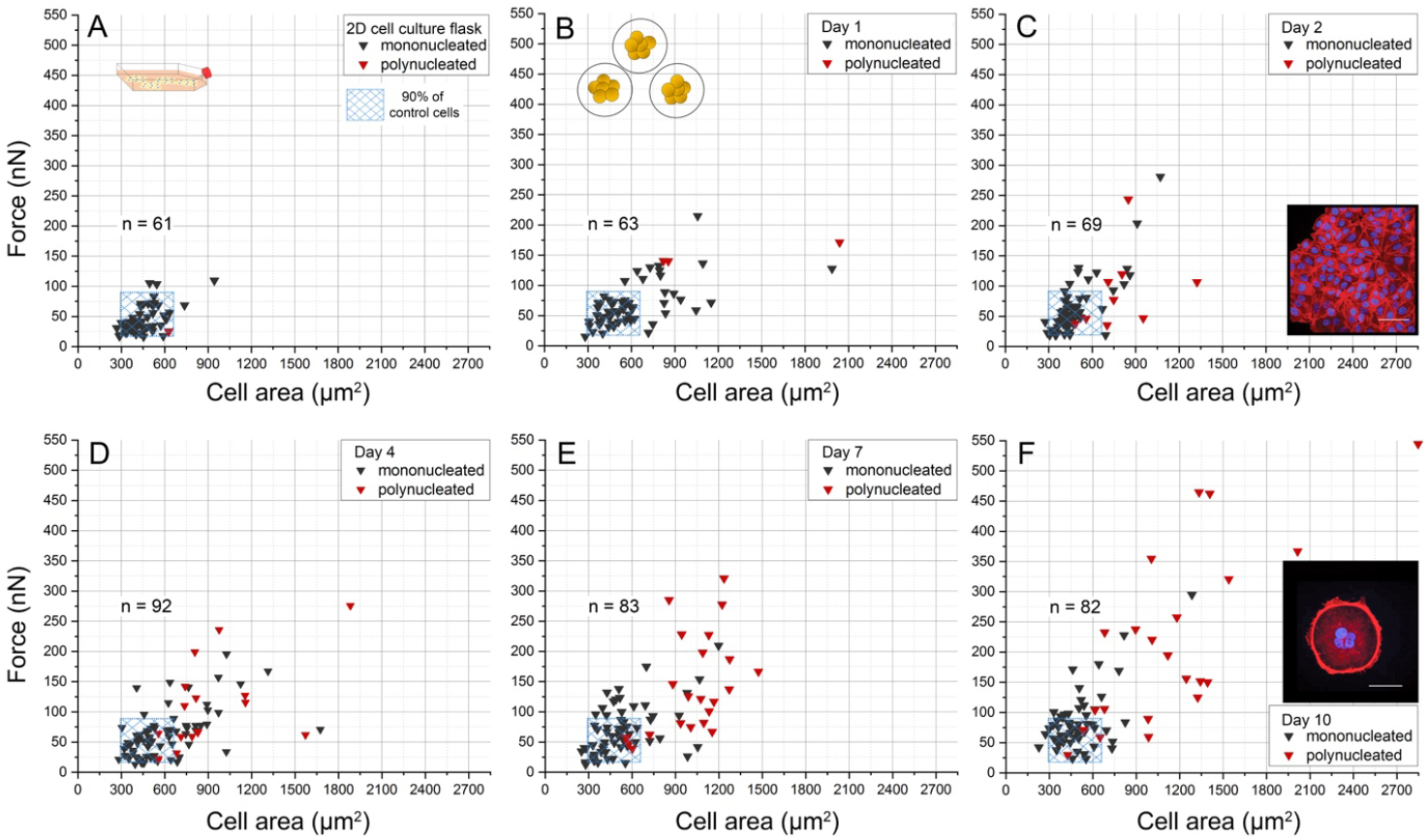
Morpho-mechanical characterization of MCF7 cells cultured on flasks (A) or those isolated from alginate-gelatin microcapsules (B-F). In the latter case, cells were isolated at days 1 (B), 2 (C), 4 (D), 7 (E) and 10 (F) post encapsulation. In all plots, the blue square represents 90% of the measurements for each dimension of the cells cultured on flasks and is used as optic reference in each graphic. In all graphics, “n” represents the number of single cells analyzed. The images located in (C) and (F) show the characteristic morphologies of cells isolated from the tumor-like microcapsules. Please note that the image placed in (F) shows a single MCF7 cell having four nuclei. In these pictures, the red staining corresponds to the actin cytoskeleton, while the blue color depicts the presence of the nuclei. The scale bar corresponds to 25 μm.

In order to compare all the assayed conditions, the magnitude of total forces exerted by cancer cells are displayed in Figures 4A and 4B. Furthermore, the F/A ratio was calculated for each condition (Figures 4C and 4D). As these Figures show, MCF7 cultured as suspended spheroids exhibit a peak of forces between days 4 and 7 post cell seeding, while cells isolated from hydrogel-based microcapsules enhance their biomechanical activity progressively. Concerning the F/A ratio, this parameter is relevant to determine if enhanced forces shown in previous figures are due to changes in their biomechanical activity, or if this is just an effect related to their increased anchorage surfaces.

**Figure 4:**
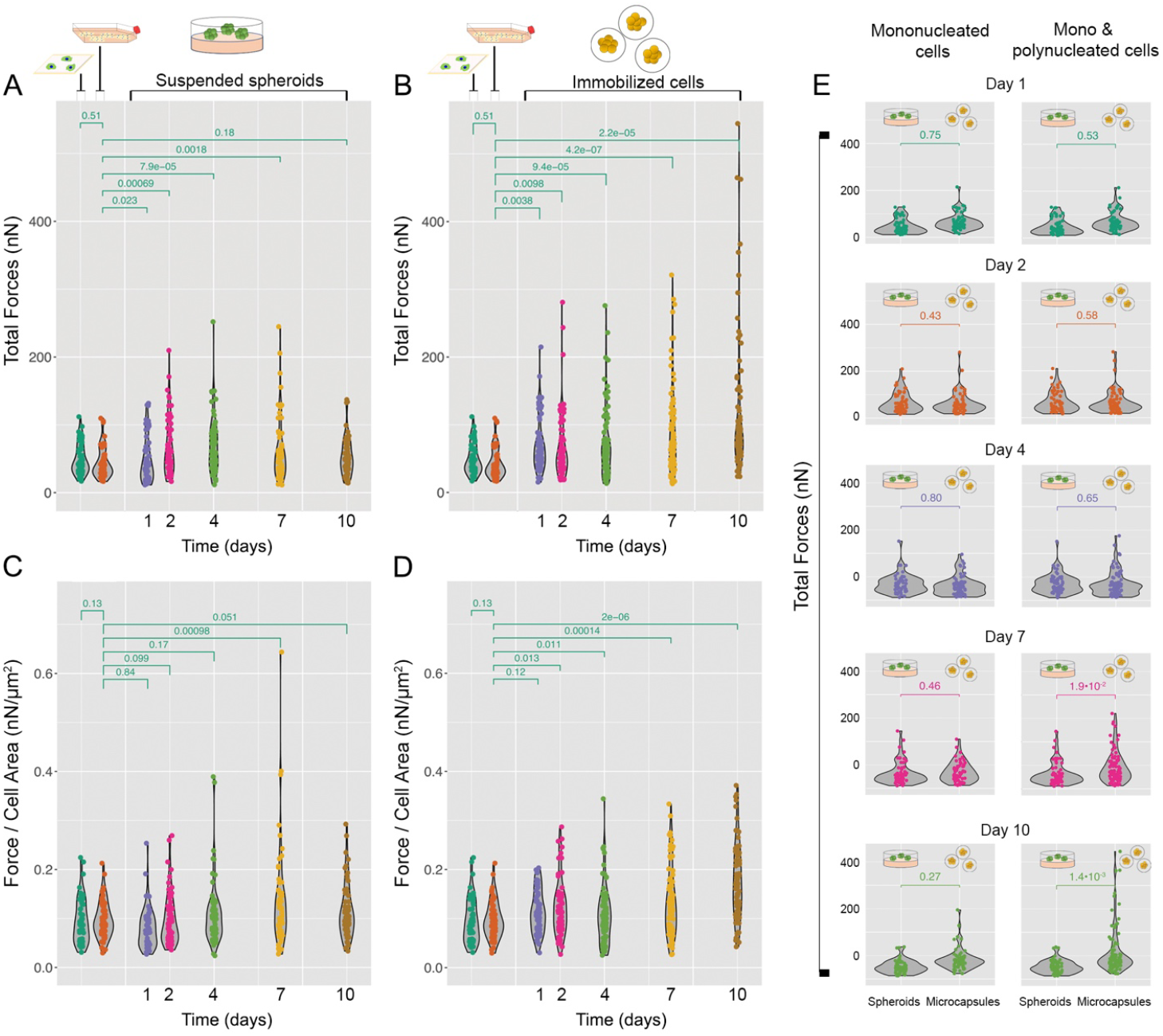
Comparative analysis of the biomechanical activity of MCF7 cells cultured in different milieus used in this work. Comparison of total forces exerted by cells isolated from spheroids (A) or from hydrogel-based microcapsules (B). Comparison of the F/A ratio between cells isolated from spheroids (C) or from hydrogel-based microcapsules (D). Forces exerted by cells isolated from spheroids or microcapsules are displayed in the panel (E). These assays were carried out with mononucleated cells (left panel), or mono- and poly-nucleated cells (right panel). In all figures, each point represents one single cell, and differences in heterogeneity were calculated using a Kolmogorov-Smirnov distribution test.

In relation to the F/A ratio, previous publications in the field have demonstrated that the magnitude of forces exerted by individual cells is proportional to their spreading area, a behavior which is highly conserved in healthy cells.^[53–56]^ However, our results are showing major changes in this parameter, mostly in MCF7 cells cultured in 3D. Interestingly, Figure 4E demonstrates that this effect is not only related to the presence of polynucleated cells, but the existence of stronger mono-nucleated cells, with respect to the population cultured on 2D (days 1, 2 and 4). Otherwise, major differences in the exertion of forces between both 3D systems are observed at days 7 and 10 post seeding/encapsulation, which seem to be indeed a result of the presence of polynucleated cells.

These results are in agreement with recent publications from Beri *et al.,* who reported the existence of at least 3 different cell sub-types of MDA-MB-231 cells, differing in their cell-matrix adhesive capabilities.^[57]^ Additionally, Yeoman *et al*. recently showed that these breast cancer cells may tune their adhesivity with time and during the progression of this disease, a pattern which may be resembled *in vitro* when entrapping these breast cancer cells in 3D milieus.^[58]^ However, none of these studies considered the presence of polyploid populations as potential responsible for that behavior, remarking the relevance of the results shown in this work.

Based on the morpho-mechanical characteristics shown by cancer cells cultured on/in different 2D and 3D milieus (Figures 1 to 4), we assumed that the mechanical stress experienced by cells cultured in 3D milieus may play a role on the generation of polyploid cells, and mostly those found in proliferative stages. To this purpose, the duplication of MCF7 was analyzed in spheroids, as well as in cells immobilized in tumor-like microcapsules (Figure 5). These assays were carried out in 3D, using methodologies already published.^[28,29,33]^

**Figure 5:**
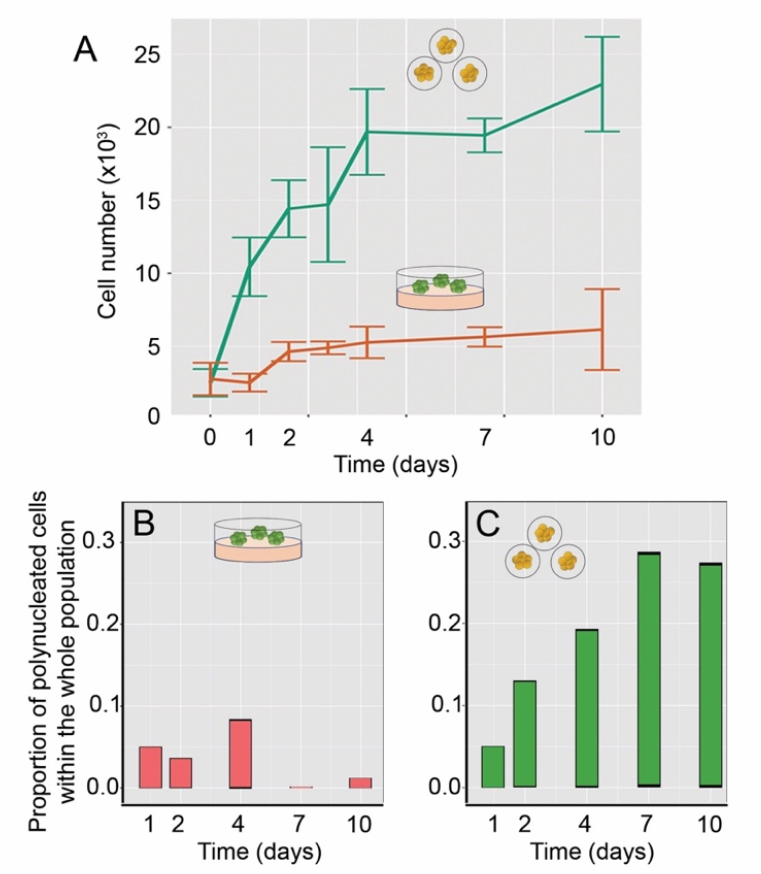
(A) Proliferation profile for cells cultured as spheroids (red line) or entrapped inside the tumorlike microcapsules (green lines). Images (B) and (C) show the proportion of polynucleated cancer cells found in the whole cell population, and isolated from spheroids (B) or from the hydrogel-based microcapsules (C).

With respect to the generation of polyploid cells, our results show that the enhanced number of multinucleated MCF7 found in the microcapsules seems to be strongly related to their doubling capabilities, as shown in Figure 5.^[59,60]^ It is relevant to mention that the low proliferation rate described in MCF7 in suspension (Figure 5A, red curve) have been previously reported, showing that this cell line exhibits a weaker duplication in absence of matrix-anchorage, compared to very pathogenic breast cancer cells (i.e. MDA-MB-231, SKBR3 or BT474).^[61]^

Based on these results, we believe that highly proliferative cells located in confined spaces, may increase the probabilities to merge their membranes, triggering to cell-cell fusion. With respect to the disrupted cytokinesis, the limited space, the stiff milieu surrounding the cancer cells, and other biological reasons (i.e., such as the restricted expression of Rho-GTPases),^[62]^ may interfere with generation of the contractile ring involved in the separation of cancer cells in duplication stages, stimulating their polyploidization. However, the cell proximity, the lack or cell-matrix anchorage, the existence of nutrient starvation or even deficiencies in the expression of E-cadherin may also lead cancer cells to become polyploid by other mechanisms, such as cell cannibalisms, entosis or emperipolesis.^[20,63]^

To test this hypothesis, cells were then characterized by RNA-seq and transcriptomic methods. Initially, we determined the presence of upregulated genes involved in polyploidization (Figure 6). To this purpose cells were isolated from spheroids and/or from polymer-based microcapsules, and upregulated genes involved in polyploidy were compared, resulting in 1832 differential expressed genes (Figures 6B and 6C). It is important to mention that the isolation of RNA was performed after 5 days post cell seeding/encapsulation. This time point was chosen, based on the lower statistical difference in terms of total forces exhibited by both cell types cultured in 3D.

**Figure 6:**
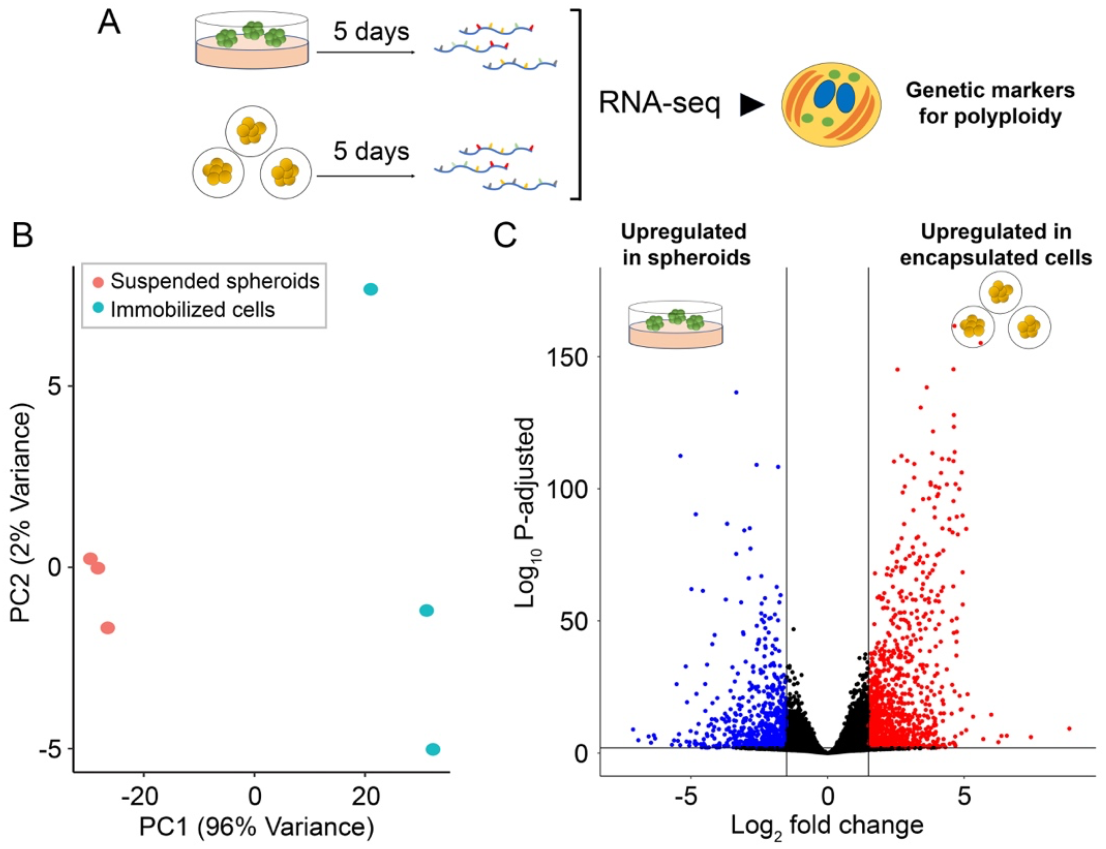
(A) Experimental design used to determine the expression of genes characterizing the presence of polynucleated cells. (B) Principal Component Analysis (PCA) of RNA-seq samples carried out with cells cultured as suspended spheroids (red dots) and cells confined in alginate–gelatin microcapsules (light-blue dots) after five days post cell seeding/immobilization (samples in triplicate). (C) The volcano plot shows the upregulated genes characterizing polyploidy in suspended spheroids (left) or in cells isolated from the tumor-like microcapsules (right) (p-adjusted < 0.01; and log2FC-1.5/1.5).

Additionally, and with the purpose to test if the populations cultured in different 3D milieus are tuning several metabolic pathways, GSEA assays were carried out (Figure 7). This image shows that cells isolated from hydrogel-based microcapsules are likely activating pathways involved in proliferation, migration and cell adhesion, supporting the results obtained by scTFM and proliferation (Figures 3, 4 and 5C respectively). It is also worthily to note the increased capability of cells extracted from the tumor-like scaffolds to repair their DNA, which can be associated to their enhanced resistance to anticancer drugs, as previously described in our last publications.^[28,29]^ On the other hand, cells isolated from spheroids are showing a lower metabolic activity, in which the autophagia and catabolic pathways are upregulated, supporting the idea that these cells would be likely activating genes involved in cell-in-cell or cell-eat-cell, instead of cell-cell fusion.

**Figure 7:**
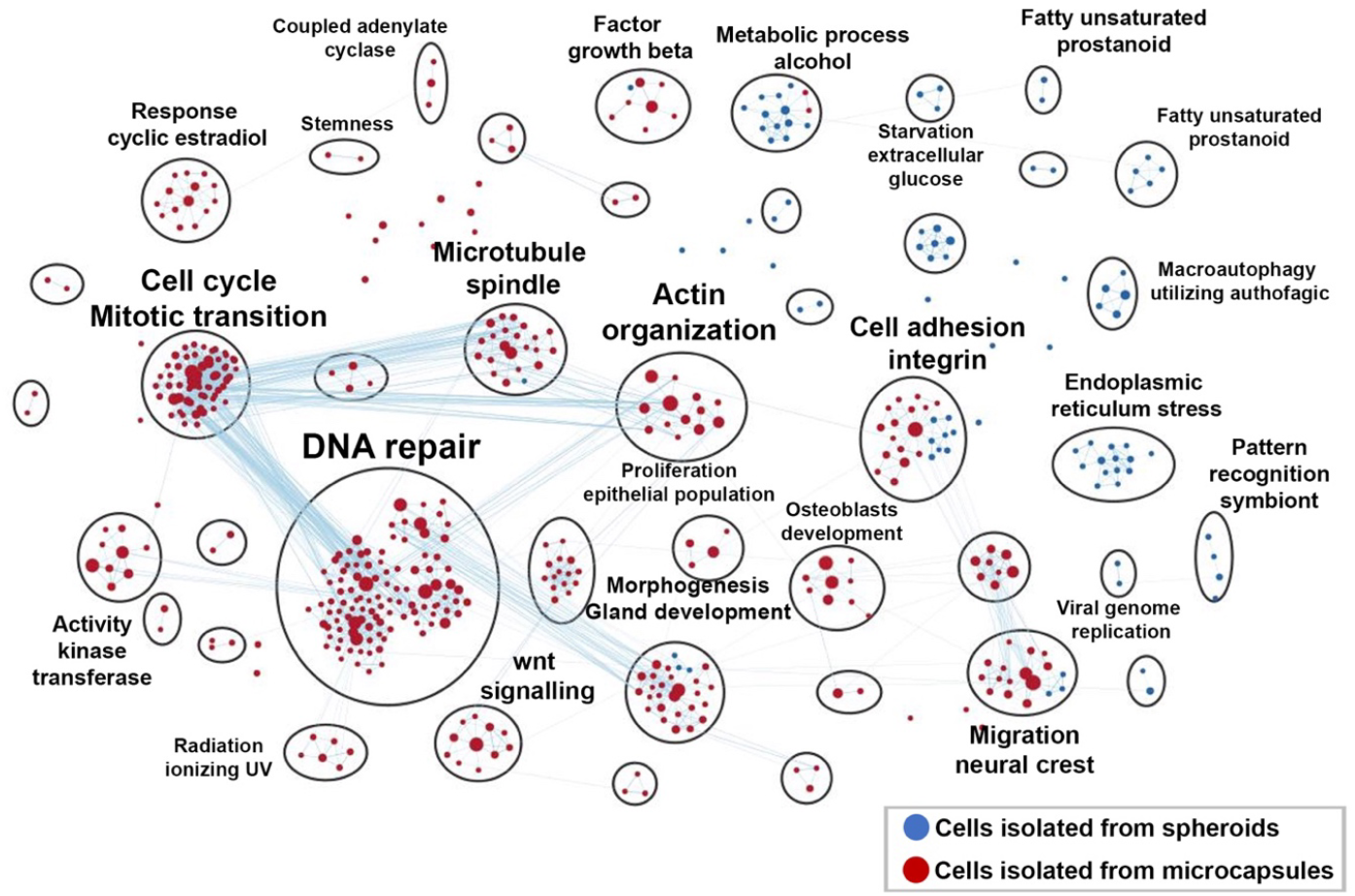
GSEA of enriched GO biological processes terms carried out in cells isolated from alginate-gelatin microcapsules and in cells cultured as spheroids (dysregulated genes, p-adjusted <0.01).

With the purpose to study this hypothesis, we performed gene expression assays, dividing the study according to the two main mechanisms responsible for cell polyploidization: i) Cell cannibalisms, entosis or emperipolesis, and ii) Cell-Cell fusion and/or disrupted cytokinesis. It is important to mention that the last two are the most described to explain the presence of Polyploid Giant Cancer Cells (PGCC) (Figure 8A).

**Figure 8:**
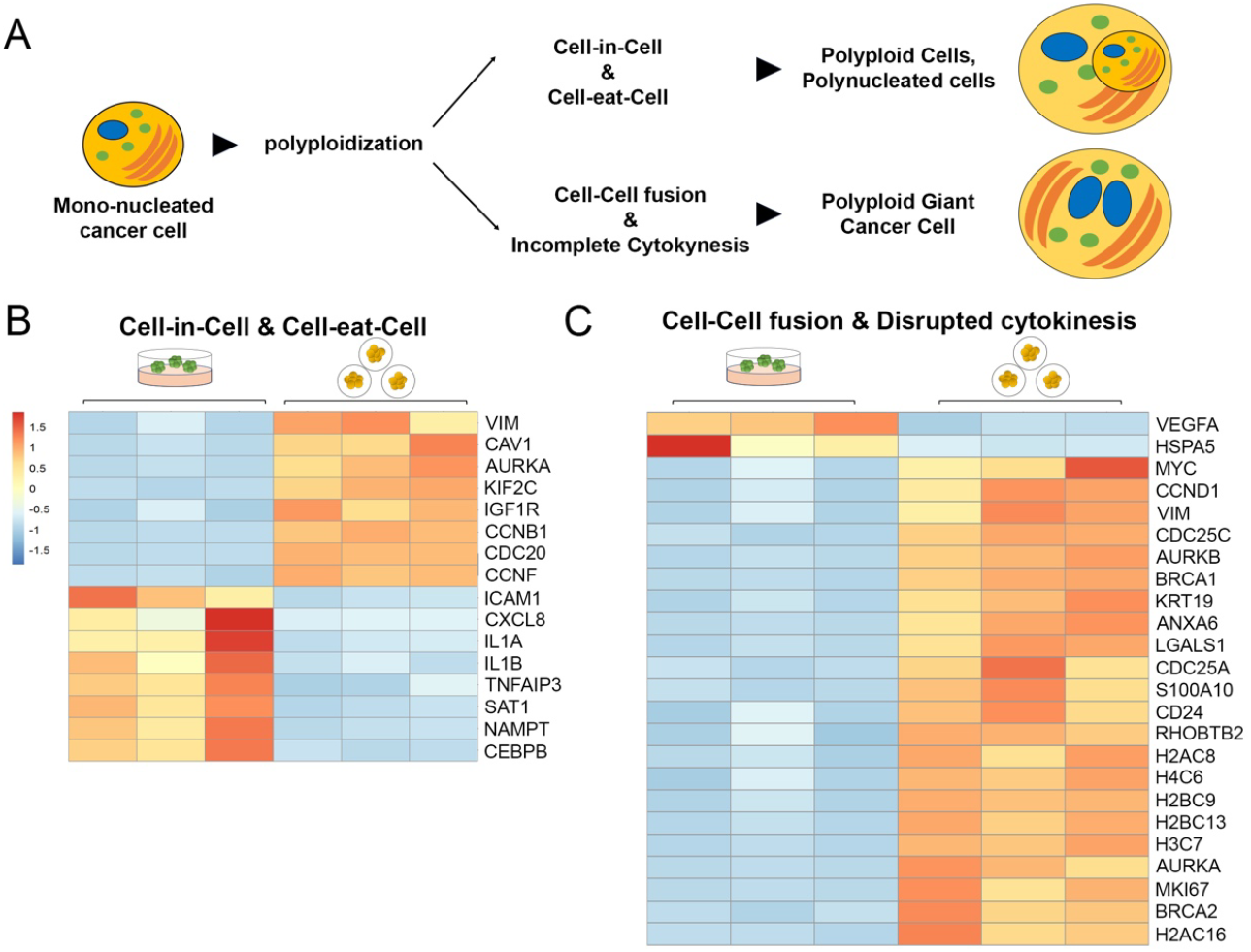
(A) Exemplification of the genetic assays carried out in this work. In this case, we performed a differential expression analysis, comparing the two main mechanisms already described in the generation polyploid cells: i) cell-in-cell and cell-eat-cell, or ii) cell-cell fusion and disrupted cytokinesis. These studies were carried out in triplicate, with cells isolated from the suspended spheroids or from the alginate-gelatin microcapsules. Heat map showing the differential expression of genes characterizing cell-in-cell and cell-eat-cell processes (B), or cell-cell fusion and disrupted cytokinesis processes (C) in breast cancer cells (p-adjusted < 0.01; log2FC −1.5/1.5).

Since most of the genes involved in each mechanism have been already reported (Supplementary Table S2), we used RNA-seq to test which strategy would be predominant in each 3D experimental condition. As Figure 8B shows, the polyploidization of cells isolated from spheroids seems to be likely generated by cell-in-cell or cell-eat-cell mechanisms, in which IL1A, IL1B, CXCL8, ICAM1, or TNFAIPR3 genes are directly involved in the process.^[14–17]^ Otherwise, cells isolated from the tumor-like microcapsules are mostly upregulating the expression of genes involved in membrane fusion *(i.e*.ANXA6, S100A10, AURKA, AURKB, LGALS1),^[64–67]^ cell proliferation and exertion of biomechanical forces (*i.e*. MKI67, KRT19, VIM, CDC25C),^[68–71]^ as well as nuclei reorganization (i.e. H2AC8, H4C6, H2BC9, H2BC13, H3C7, H2AC16),^[72,73]^ suggesting that the 3D polymer-based scaffolds may restrict -but not hinder-their proliferation, facilitating the cell-cell fusion and/or interfering with the cellular cytokinesis, enabling the generation of PGCC. However, it is important to mention that cells isolated from the tumor-like microcapsules are also upregulating the expression of specific markers for cell cannibalism (i.e. VIM, AURKA),^[20,74–77]^ suggesting that polyploidization of confined cells located in the hydrogel-based scaffolds may be also triggered by these mechanisms.

It is finally relevant to comment that although this hypothesis has not been properly tested in this work, we can envisage alternatives to study by which mechanism polyploid cells are generated, such as by Karyotyping studies ^[78]^ combined with imaging assays.^[79]^ Yet, our outcomes are indicating that MCF7 cells need both the confinement in a 3D milieu and the mechanical stress to generate polynucleated cells, task which can be easily carried out by immobilizing cancer cells in alginate-gelatin microcapsules.

## 4. Conclusions

Due to the biophysical relevance of cell heterogeneity in cancer progression, we focused this study on determining how the morpho-mechanical properties of cancer cells may be influenced by the milieu where they are cultured, and how polynucleated cells are generated under different culture conditions. For this purpose, the breast cancer cell line MCF7 was cultured in different 2D and 3D scaffolds, enabling their preconditioning prior characterizing them by biophysical and transcriptomic methods. Special attention was paid on the determination of a special type of polyploid cells, known as Polyploid Giant Cancer Cells, due to their relevance in cancer progression. As already shown, the increased presence of PGCC in the tumor-like microcapsules strongly suggests that they are likely generated during their proliferation in confined environments, and under mechanical stress. Remarkably, these results also show that no other physical, chemical, or biological stimuli are required to generate them. Thus, and as a major result, this work demonstrate that polymer-based strategies may offer the scientific community a new alternative for generating polyploid cells in a system much more akin to *in vivo* tumors.

Finally, our results also note that the time used to pre-condition cancer cells may also play a relevant role in the apparition of stronger mononucleated populations, as well as polynucleated cells. These outcomes should encourage the scientific community to be aware about the time-point used to obtain biological samples from *in vitro* systems, in order to perform reliable and comparable analysis of cancer malignancy, including but not restricted to, adhesion and migration studies, as well as cytotoxicological tests to name a couple of examples.

## Data Availability Statement

The data that support the findings of this study are available from the corresponding author upon reasonable request.

## Acknowledgments

Aldo Leal-Egaña acknowledges the financial support provided by the German Research Foundation (LE3418/4-1 and LE3418/2-1). Anja Bosserhoff acknowledges the support by the German Research Foundation (TRR225, project C03). Miguel Fuentes-Chandía thanks CONICYT for his postdoctoral fellowship. All authors gratefully thank Alexander-Oliver Matthies for preparation of RNA samples and RNA-Seq libraries, Dr. Trevor Kalkus and Dr. Philipp Trippal for constructive discussions during the preparation of this work, and the Biomedical Sequencing Facility (BSF) of the CeMM (Vienna, Austria) for sequencing our samples. Johannes Schödel and René Krüger were supported by the Deutsche Forschungsgemeinschaft (DFG, German Research Foundation), project number 387509280, SFB 1350 (project C5). Finally, the use of the spinning disc confocal microscope (Zeiss Spinning Disc Axio Observer Z1) was possible due to the funded project 248122450, from German Research Foundation (DFG).

---

**Figure S1:**
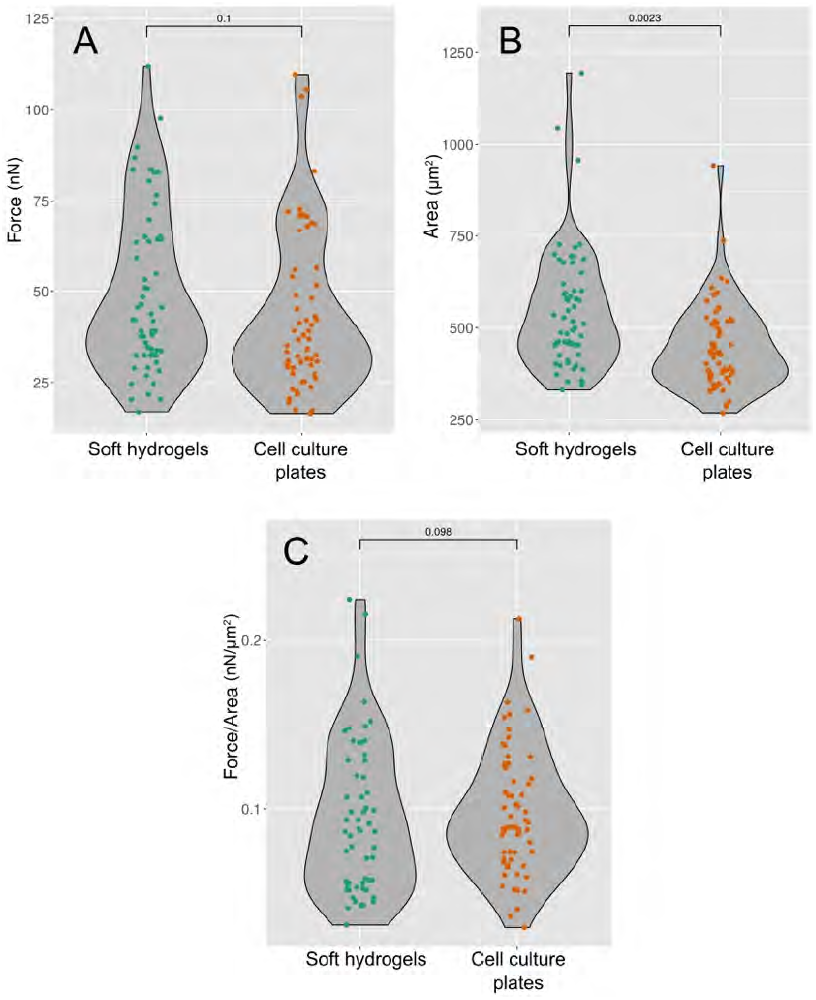
Comparative analysis of forces (A), spreading area (B), and the Force/Area ratio (C) between cells cultured on soft hydrogels, or on cell culture plates (control experiments). In these assays, MCF7 cells were analyzed after 5 days post cell seeding. Statistical differences were calculated using a Kolmogorov-Smirnov distribution test.

**Supplementary Table 2A:**
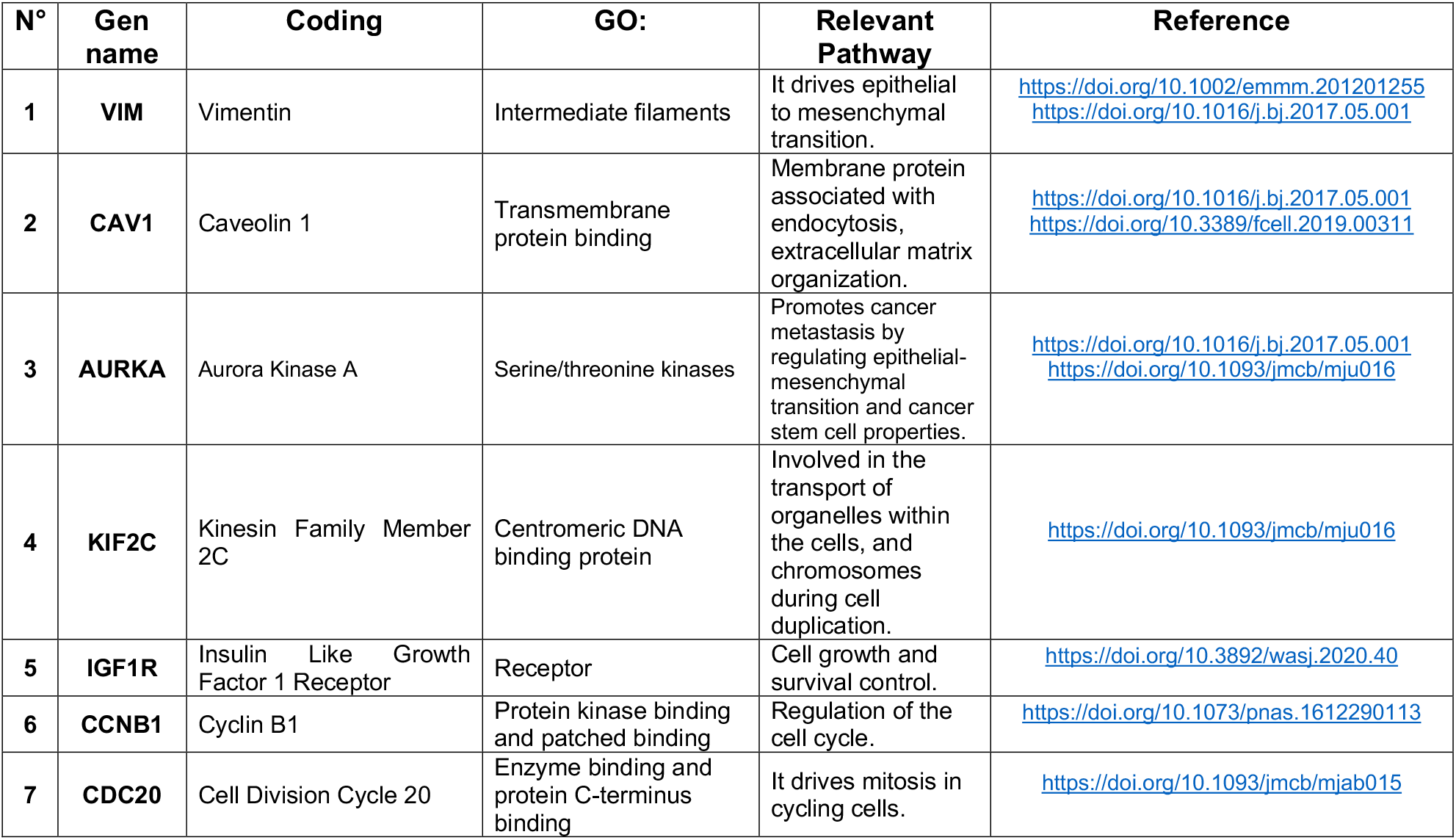
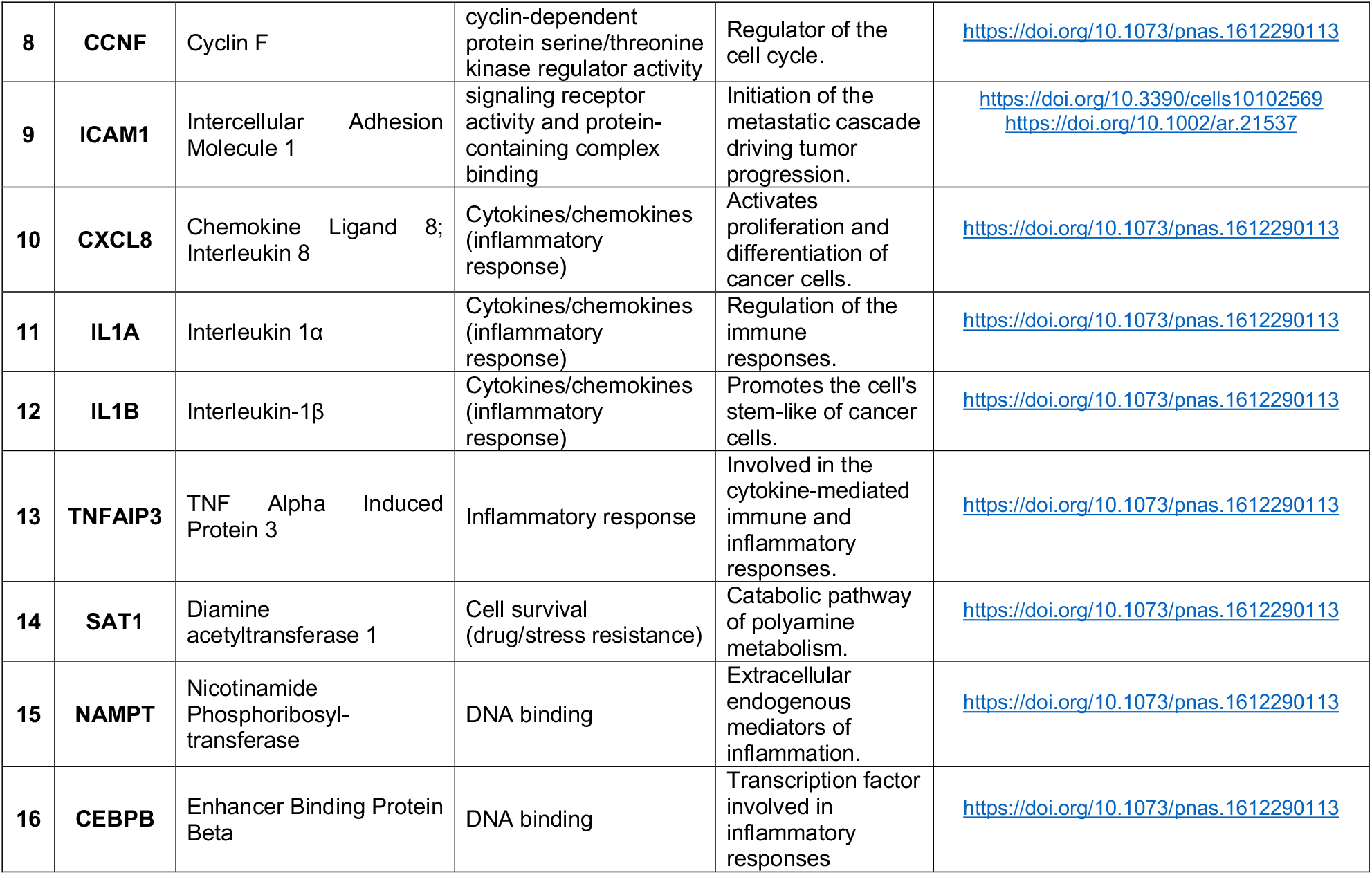
Genes characterizing cell-in-cell or cell-eat-cell processes

| **N°** | **Gen name** | **Coding** | **GO:** | **Relevant Pathway** | **Reference** |
| --- | --- | --- | --- | --- | --- |
| 1 | **VIM** | Vimentin | Intermediate filaments | It drives epithelial to mesenchymal transition. | https://doi.org/10.1002/emmm.201201255<br>https://doi.org/10.1016/j.bj.2017.05.001 |
| 2 | **CAV1** | Caveolin 1 | Transmembrane protein binding | Membrane protein associated with endocytosis, extracellular matrix organization. | https://doi.org/10.1016/j.bj.2017.05.001<br>https://doi.org/10.3389/fcell.2019.00311 |
| 3 | **AURKA** | Aurora Kinase A | Serine/threonine kinases | Promotes cancer metastasis by regulating epithelial-mesenchymal transition and cancer stem cell properties. | https://doi.org/10.1016/j.bj.2017.05.001<br>https://doi.org/10.1093/jmcb/mju016 |
| 4 | **KIF2C** | Kinesin Family Member 2C | Centromeric DNA binding protein | Involved in the transport of organelles within the cells, and chromosomes during cell duplication. | https://doi.org/10.1093/jmcb/mju016 |
| 5 | **IGF1R** | Insulin Like Growth Factor 1 Receptor | Receptor | Cell growth and survival control. | https://doi.org/10.3892/wasj.2020.40 |
| 6 | **CCNB1** | Cyclin B1 | Protein kinase binding and patched binding | Regulation of the cell cycle. | https://doi.org/10.1073/pnas.1612290113 |
| 7 | **CDC20** | Cell Division Cycle 20 | Enzyme binding and protein C-terminus binding | It drives mitosis in cycling cells. | https://doi.org/10.1093/jmcb/mjab015 |
| 8 | **CCNF** | Cyclin F | cyclin-dependent protein serine/threonine kinase regulator activity | Regulator of the cell cycle. | https://doi.org/10.1073/pnas.1612290113 |
| 9 | **ICAM1** | Intercellular Adhesion Molecule 1 | signaling receptor activity and protein-containing complex binding | Initiation of the metastatic cascade driving tumor progression. | https://doi.org/10.3390/cells10102569<br>https://doi.org/10.1002/ar.21537 |
| 10 | **CXCL8** | Chemokine Ligand 8; Interleukin 8 | Cytokines/chemokines (inflammatory response) | Activates proliferation and differentiation of cancer cells. | https://doi.org/10.1073/pnas.1612290113 |
| 11 | **IL1A** | Interleukin 1α | Cytokines/chemokines (inflammatory response) | Regulation of the immune responses. | https://doi.org/10.1073/pnas.1612290113 |
| 12 | **IL1B** | Interleukin-1β | Cytokines/chemokines (inflammatory response) | Promotes the cell's stem-like of cancer cells. | https://doi.org/10.1073/pnas.1612290113 |
| 13 | **TNFAIP3** | TNF Alpha Induced Protein 3 | Inflammatory response | Involved in the cytokine-mediated immune and inflammatory responses. | https://doi.org/10.1073/pnas.1612290113 |
| 14 | **SAT1** | Diamine acetyltransferase 1 | Cell survival (drug/stress resistance) | Catabolic pathway of polyamine metabolism. | https://doi.org/10.1073/pnas.1612290113 |
| 15 | **NAMPT** | Nicotinamide Phosphoribosyl-transferase | DNA binding | Extracellular endogenous mediators of inflammation. | https://doi.org/10.1073/pnas.1612290113 |
| 16 | **CEBPB** | Enhancer Binding Protein Beta | DNA binding | Transcription factor involved in inflammatory responses | https://doi.org/10.1073/pnas.1612290113 |

**Supplementary Table 2B:**
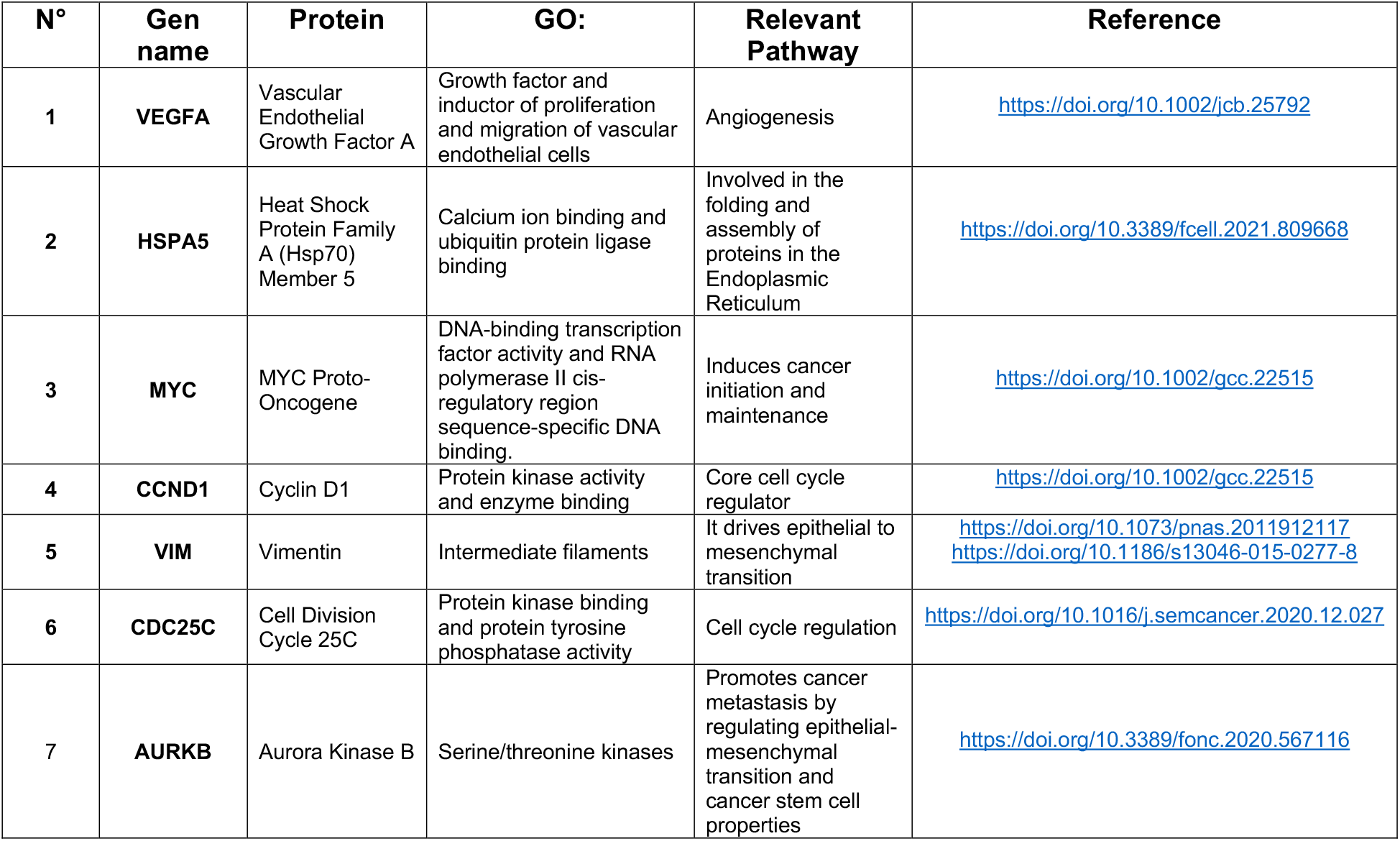
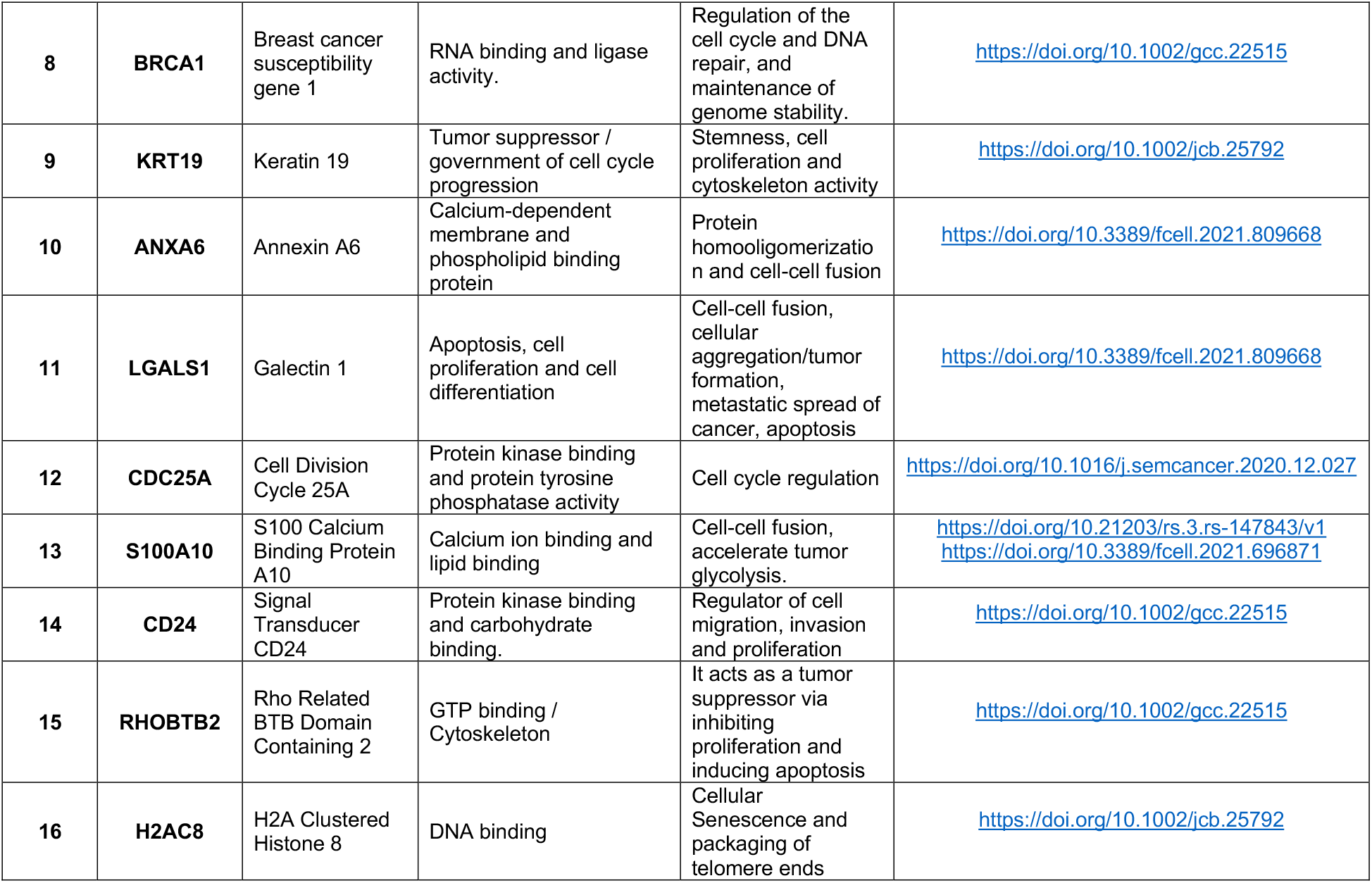
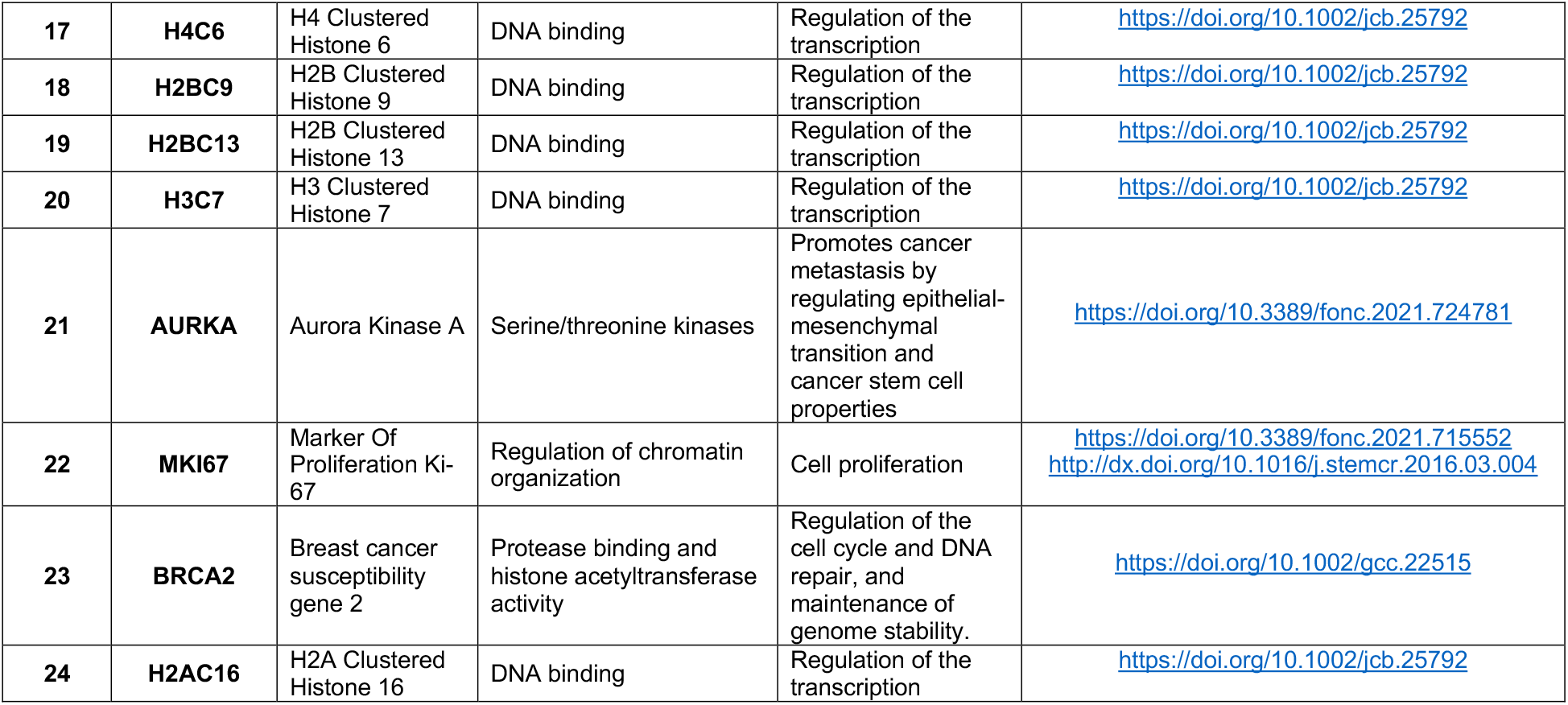
Genes characterizing cell-cell fusion or incomplete cytokinesis.

